# NGScloud2: optimized bioinformatic analysis using Amazon Web Services

**DOI:** 10.1101/2020.11.30.400010

**Authors:** Fernando Mora-Márquez, José Luis Vázquez-Poletti, Unai López de Heredia

## Abstract

NGScloud was a bioinformatic system developed to perform *de novo* RNAseq analysis of non-model species by exploiting the cloud computing capabilities of Amazon Web Services. The rapid changes undergone in the way this cloud computing service operates, along with the continuous release of novel bioinformatic applications to analyze next generation sequencing data, have made the software obsolete. NGScloud2 is an enhanced and expanded version of NGScloud that permits the access to *ad hoc* cloud computing infrastructure, scaled according to the complexity of each experiment. NGScloud2 presents major technical improvements, such as the possibility of running spot instances and the most updated AWS instances types, that can lead to significant cost savings. As compared to its initial implementation, this improved version updates and includes common applications for *de novo* RNAseq analysis, and incorporates tools to operate workflows of bioinformatic analysis of reference-based RNAseq, RADseq and functional annotation. NGScloud2 optimizes the access to Amazon’s large computing infrastructures to easily run popular bioinformatic software applications, otherwise inaccessible to non-specialized users lacking suitable hardware infrastructures. The correct performance of the pipelines for *de novo* RNAseq, reference-based RNAseq, RADseq and functional annotation was tested with real experimental data. NGScloud2 code, instructions for software installation and use are available at https://github.com/GGFHF/NGScloud2. NGScloud2 includes a companion package, NGShelper that contains python utilities to post-process the output of the pipelines for downstream analysis at https://github.com/GGFHF/NGShelper.

## Introduction

The large output size of Next Generation Sequencing (NGS) technologies and the algorithms and applications employed in their analysis, present processing limitations typical of big data, such as RAM size, CPU capacity, storage and data accessibility (Yang et al., 2017). Therefore, research labs have to allocate a significant part of their budget to provisioning, managing and maintaining their computational infrastructure (Kwon et al., 2015). A cost-efficient alternative for NGS analysis that presents several advantages over local or HPC hardware infrastructure resides in cloud computing (Langmead & Nellore, 2018). Cloud computing is flexible and scalable, allowing various configurations of OS, RAM size, CPU number and almost unlimited storage to fit the hardware resources for a specific bioinformatic workflow. Once the workflow computing requirements are provisioned, hardware resources are readily available, and the workflow performance and data can be securely accessed and monitored at any time from any local computer with internet access. Moreover, for public cloud services, the user only pays for the effectively used resources, reducing experiment times and costs.

Here we present NGScloud2, a new version of the NGScloud software (Mora-Márquez, Vázquez-Poletti & López de Heredia U, 2018). NGScloud was developed as a bioinformatic system to perform *de novo* RNAseq analysis of non-model species. This was accomplished using the cloud computing infrastructure from Amazon Web Services (AWS), the Elastic Compute Cloud (EC2), and its high-performance block storage service, the Amazon Elastic Block Store (EBS). NGScloud allowed to create one or more EC2 instances (virtual machines) of M3, C3 or R3 instance types forming clusters where analytic processes were run using StarCluster, an open source clustercomputing toolkit for EC2 (http://star.mit.edu/cluster/). However, NGScloud did not support the new instance types that AWS has made available since the original application release. Below we describe the major new features of NGScloud2 that significantly expand NGScloud2 functionality with respect to the original version.

## Materials & Methods

NGScloud2 is a free and open source program written in Python3. Source code and a complete manual with installation instructions and tutorials to exploit all the potential of NGScloud2 are available from the GitHub repository (https://github.com/GGFHF/NGScloud2). NGScloud2 presents remarkable differences with respect to NGScloud both in the way AWS resources are managed to better exploit all the potential of EC2 and EBS, but also by incorporating the possibility of running a more complete set of bioinformatic applications and pipelines for *de novo* RNAseq, reference-based RNAseq, Restriction site Associated DNA sequencing (RADseq) and functional annotation (see Results and Discussion section). In addition, a toolkit of Python programs useful to post-process the output of RNAseq and RADseq experiments is available in NGShelper (https://github.com/GGFHF/NGShelper).

The correct operability of the pipelines for *de novo* RNAseq, reference-based RNAseq, RADseq and funcional annotation was tested with data generated by our research group. Test data for RNAseq and RADseq workflows consisted of two sets of Illumina^TM^ reads: (1) Pcan, a paired-ended RNA library of xylem regeneration tissue of the conifer tree *Pinus canariensis* (Mora-Márquez et al. 2020a). (2) Suberintro, a set of 16 paired ended Illumina™ libraries of *Quercus suber, Quercus ilex* and their hybrids obtained from leaf tissue; eight libraries correspond to genotyping-by-sequencing with MslI and other eight libraries correspond to ddRADseq with PstI-MspI (see details in Guillardín-Calvo et al., 2019). Read data are available at NCBI: SRX5228139 – SRX5228161 for Pcan, and SRX5019123-SRX5019138 for Suberintro. The functional annotation workflow was tested with a small subset of transcripts corresponding to the monolignol biosynthesis gene family in Arabidopsis (Raes et al., 2003).

## Results & Discussion

### Technical improvements

NGScloud2 introduces a more efficient architecture of instances and volumes than the original version (Figure 1). While NGScloud used one volume for each type of existing datasets (applications, databases, references, reads and results), NGScloud2 offers the possibility of holding all dataset types in a unique volume, thus reducing the complexity in volume management. NGSCloud2 philosophy is based on the “cluster” concept. A cluster is a set of 1 to n virtual machines with the same instance type. Each instance type has its hardware features: processor type, CPU number, memory amount, etc. (https://aws.amazon.com/ec2/instance-types/).

**Figure 1.**
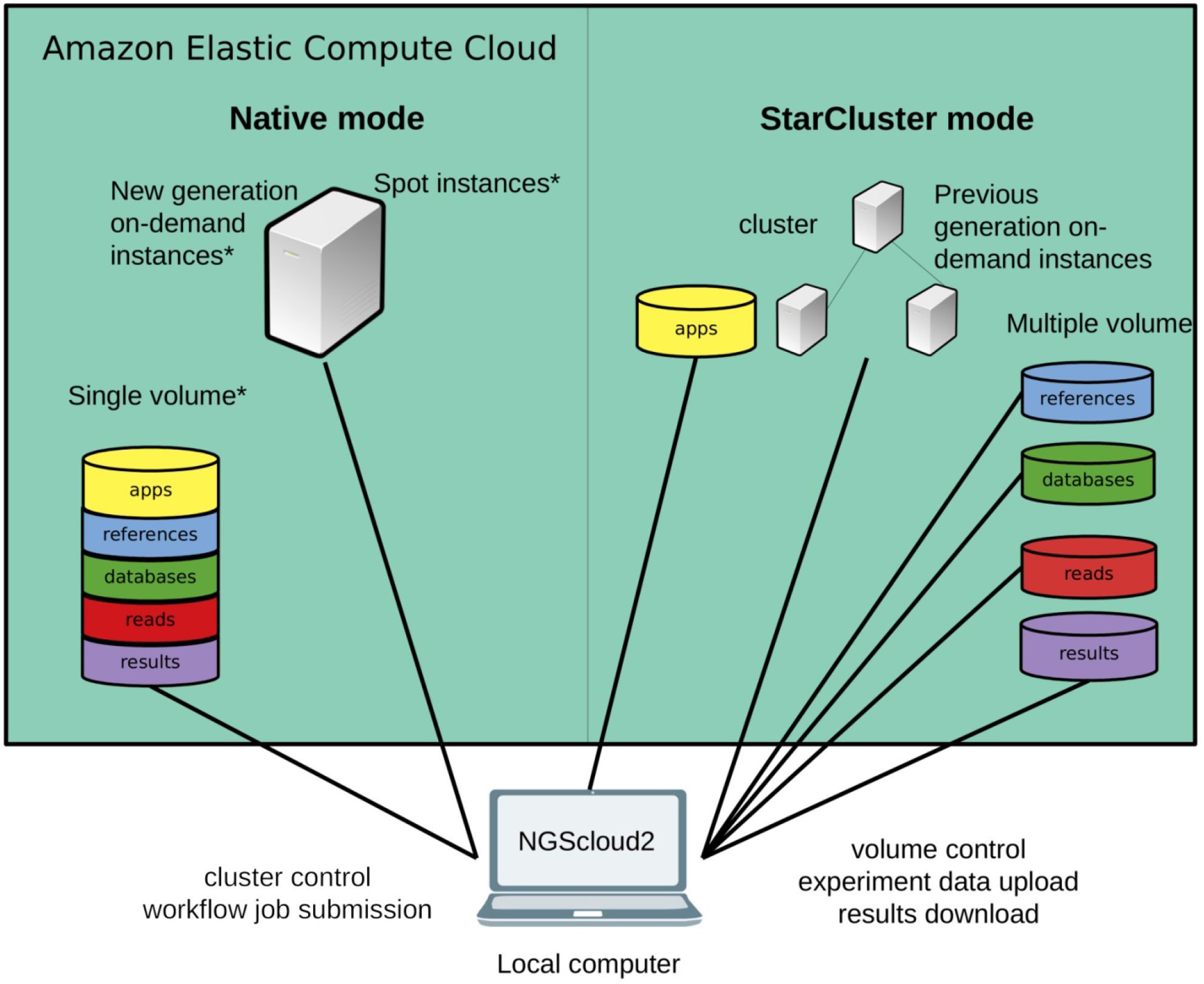
Technical improvements of NGScloud2.

NGScloud2 includes two cluster modes, StarCluster and native. The StarCluster mode uses StarCluster (http://star.mit.edu/cluster/), an open source cluster-computing toolkit for EC2, which implements clusters of up to 20 virtual machines, enabling faster analysis. The last version of Starcluster (0.95.6) dates from 2013 and can only use AWS’s previous generation instance types, i.e. m3, c3 or r3. In NGScloud2, we provide a patch to enable using m4, c4 and r4 instance types.

To reduce the dependency of NGScloud from StarCluster, which only allows to create clusters of previous generation instances, NGScloud2 has incorporated a “native” instance creation mode that sets a single virtual machine with any of the currently available on-demand EC2 instance types (m4, c4, r4, m5, m5a, c5, c5a, r5 and r5a). The new generation instance types are slightly cheaper and their hardware improves over equivalent hardware from previous generations. Moreover, the new version enables launching “spot instances” that derive from unused EC2 capacity in the AWS cloud (https://aws.amazon.com/ec2/spot/). Spot instances have the advantage of being up to 50-80% cheaper than on-demand instances at the cost of suffering unpredictable interruption out of control of the user (Supplemental Table 1). Therefore, using spot instances is highly recommended for data transfer and for certain bioinformatic processes that run fast, process small volume input or include the possibility to be re-launched from the process interruption point.

NGScloud2 includes a user-friendly graphical front-end to operate the hardware resources, submit processes, and manage the data. The front-end includes a drop-down menu to configure AWS resources (clusters, nodes and volumes) and to install available bioinformatic software. Data transfer between the cloud and the local computer is operated through another drop-down menu. Additional drop-down menus are available to run de novo RNAseq, reference-based RNAseq, RADseq and functional annotation workflows, respectively. Log files of each executed process can be consulted in the “Logs” menu.

### New methods and applications available

The other major improvements of NGScloud2 over NGScloud are related to the implementation of new bioinformatic pipelines and application tools (Table 1) that are automatically installed using Bioconda (Grüning et al., 2018), thus giving access to updated versions of the software without worrying about dependencies and software requirements. While the original purpose of NGScloud was to help in *de novo* RNAseq analysis, NGScloud2 includes pipelines and applications to perform reference based RNAseq, RADseq and functional annotation. A summary of the AWS instances employed and the total elapsed times for the pipelines run on the test data is available in Supplemental Table 2, Table3, Table 4 and Table 5.

**Table 1:**
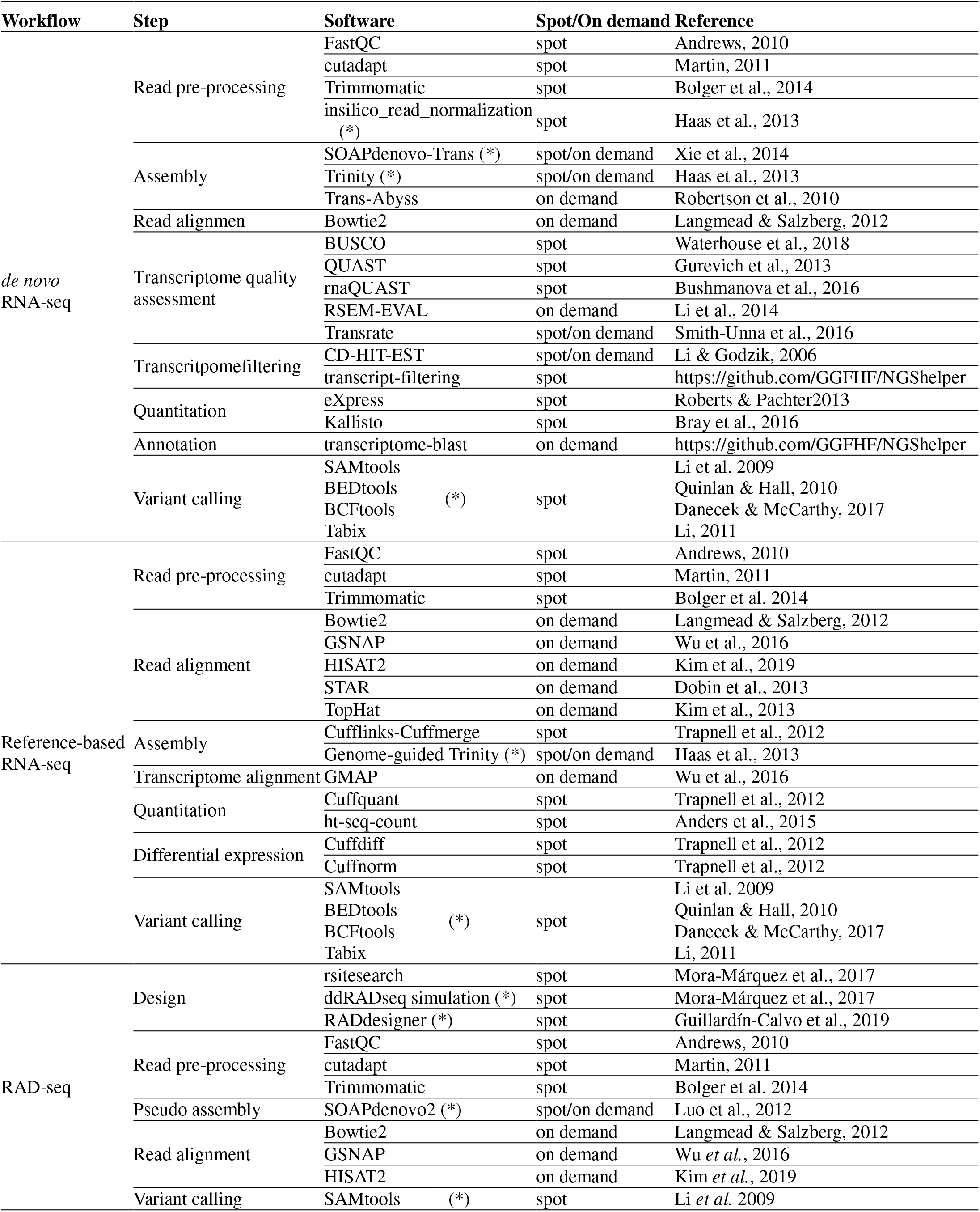
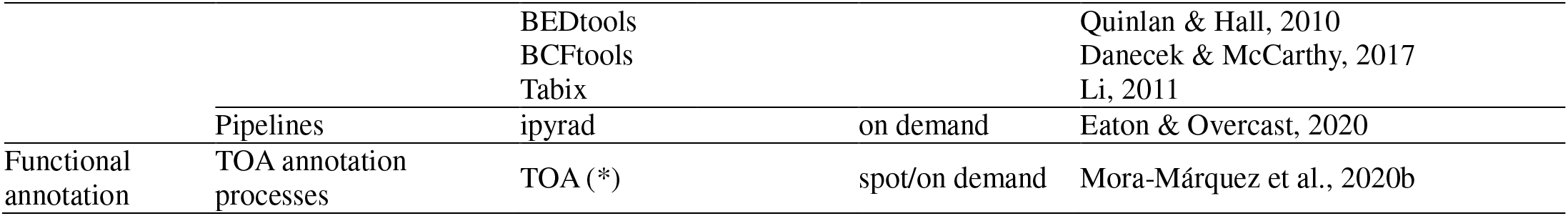
Software available for de novo RNA-seq, reference-based RNAseq, RADseq and functional annotation in NGScloud2. Recommendations for use spot or on demand instances are provided to optimize costs at every step of the workflows. (*) For time consuming processes that can be re-launched from the point of interruption, both spot or on demand instances may produce optimal performance, depending on the user’s needs.

#### *De novo* RNAseq

The original software was mainly focused on *de novo* assembly of RNAseq libraries using either Trinity, and included pre-processing of reads with FASTQC (Andrews, 2010), Trimmomatic (Bolger, Lohse & Usadel, 2014) and three *de novo* RNAseq assemblers: Trinity (Haas et al., 2013), SoapDeNovo-Trans (Xie et al., 2014) and Transabbys (Robertson et al., 2010). NGScloud2 *de novo* RNAseq workflow has been improved (Figure 2) by including cutadapt (Martin, 2011) to perform read pre-processing, a new read alignment step with Bowtie2 (Langmead & Salzberg, 2012) to map back the reads to the assembled transcriptome and software to quantify total counts of transcripts for further differential expression analysis: eXpress (Roberts & Pachter, 2013) and Kallisto (Bray el al., 2016). Intensive processes, such as Trinity and SOAPdenovo-Trans transcriptome assemblers can now be re-launched from the point where the process interruption occurred, thus preventing unexpected malfunctioning of the cloud system or software bugs (Mora-Márquez et al. 2020a). A variant calling step is also included to find SNPs or indels using SAMtools (Li et al. 2009), BEDtools (Quinlan & Hall, 2010) and BCFtools (Danecek & McCarthy, 2017).

**Figure 2.**
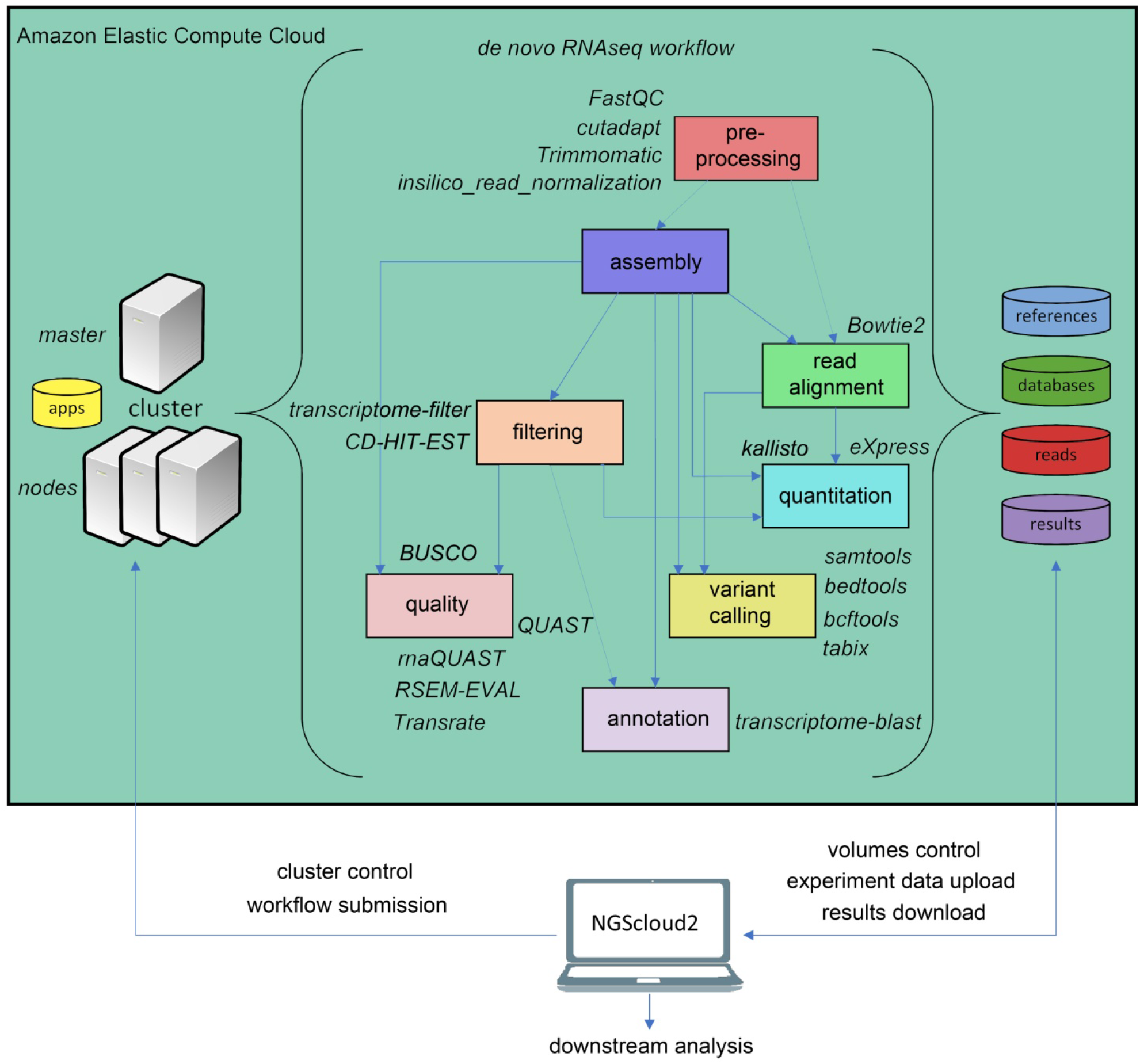
De novo RNAseq workflow in NGScloud2.

#### Reference-based RNAseq

In the last years, an increasing number of genomic and transcriptomic resources are available for many plant and animal species. Therefore, reference-based RNAseq is expected to become a usual practice not only for model species. NGScloud2 includes a workflow to accomplish read preprocessing, read alignment, reference-guided assembly, quantitation, differential expression and variant calling (Figure 3). Read pre-processing is done with the same tools as for *de novo* RNAseq (Trimmomatic and cutadapt). Read alignment to a reference genome assembly can be performed with Bowtie2, or with popular splice-aware aligners: Hisat2 (Kim et al., 2019), TopHat2 (Kim et al., 2013), STAR (Dobin et al., 2013) or GSNAP (Wu et al., 2016). Moreover, read alignment can also be run against a reference transcriptome using GMAP (Wu et al., 2016). After read alignment, a transcriptome can be assembled using Cufflinks-Cuffmerge (Trapnell et al., 2012). Reference-guided *de novo* assembly can also be performed with Trinity’s genome guided version (Haas et al., 2013). Transcript or isoform abundance can be quantified with Cuffquant (Trapnell et al., 2012) or HT-seq-count (Anders, Pyl & Huber, 2015), and differential expression analysis can be run with Cuffdiff and Cuffnorm (Trapnell et al., 2012). A variant calling step that operates in a similar way than for de novo RNA-seq is also included.

**Figure 3.**
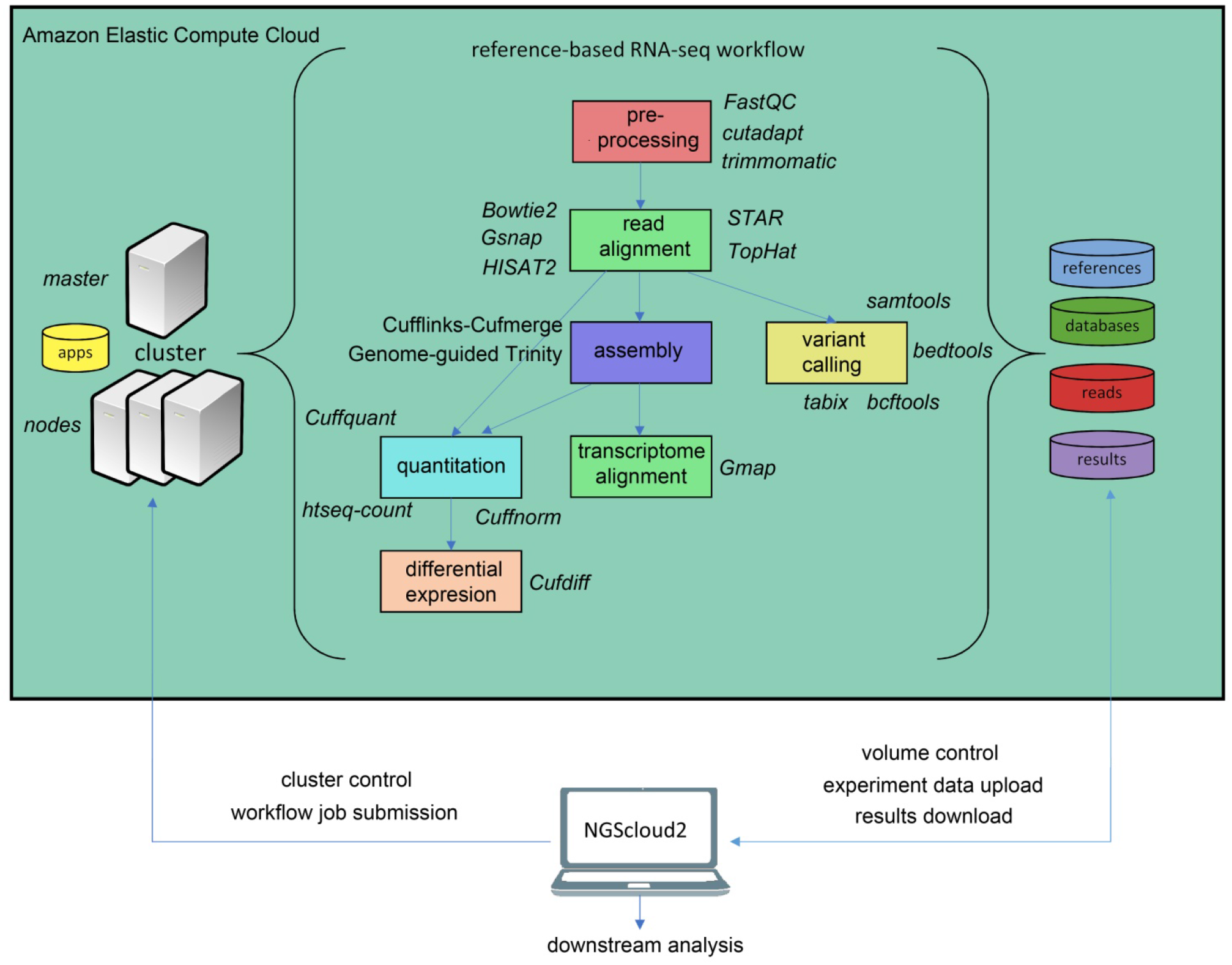
Reference-based RNAseq workflow in NGScloud2.

#### RADseq

Another major novelty in NGScloud2 is the possibility of running RAD-seq bioinformatic workflows. This reduced genome representation methodology and its derivates (e.g. ddRADseq) are used to find out polymorphism in specific genomic regions nearby restriction enzyme cut sites in populations of multiple individuals, and has revealed powerful in phylogenetics, population genetics, and association mapping studies, among others (Andrews et al., 2016). In NGScloud2, we have included ddRADseqTools (Mora-Márquez et al., 2017) and RADdesigner (Guillardín-Calvo et al., 2019) to assess the optimal experimental design of a RADseq experiment, i.e. to choose the enzyme combinations, simulate the effect of allele dropout and PCR duplicates on coverage, quantify genotyping errors, optimize polymorphism detection parameters or determine sequencing depth coverage.

The workflow of RADseq data in NGScloud2 allows to analyze the data using two strategies (Figure 4). RADseq libraries can be mapped with Bowtie2, GSNAP or HISAT2 to an available genome or psedudogenome assembly. The pseudogenome can be assembled using the same (or complementary) reads with SOAPdenovo2 genomic assembler (Luo et al., 2012), or with the Starcode sequence clusterizer (Zorita, Cuscó & Filion, 2015). After read mapping, variant calling is performed in a similar way than for *de novo* RNA-seq. The alternative is to perform read clusterization, filtering and variant calling in a single step with the robust iPyrad pipeline (Eaton & Overcast, 2020).

**Figure 4.**
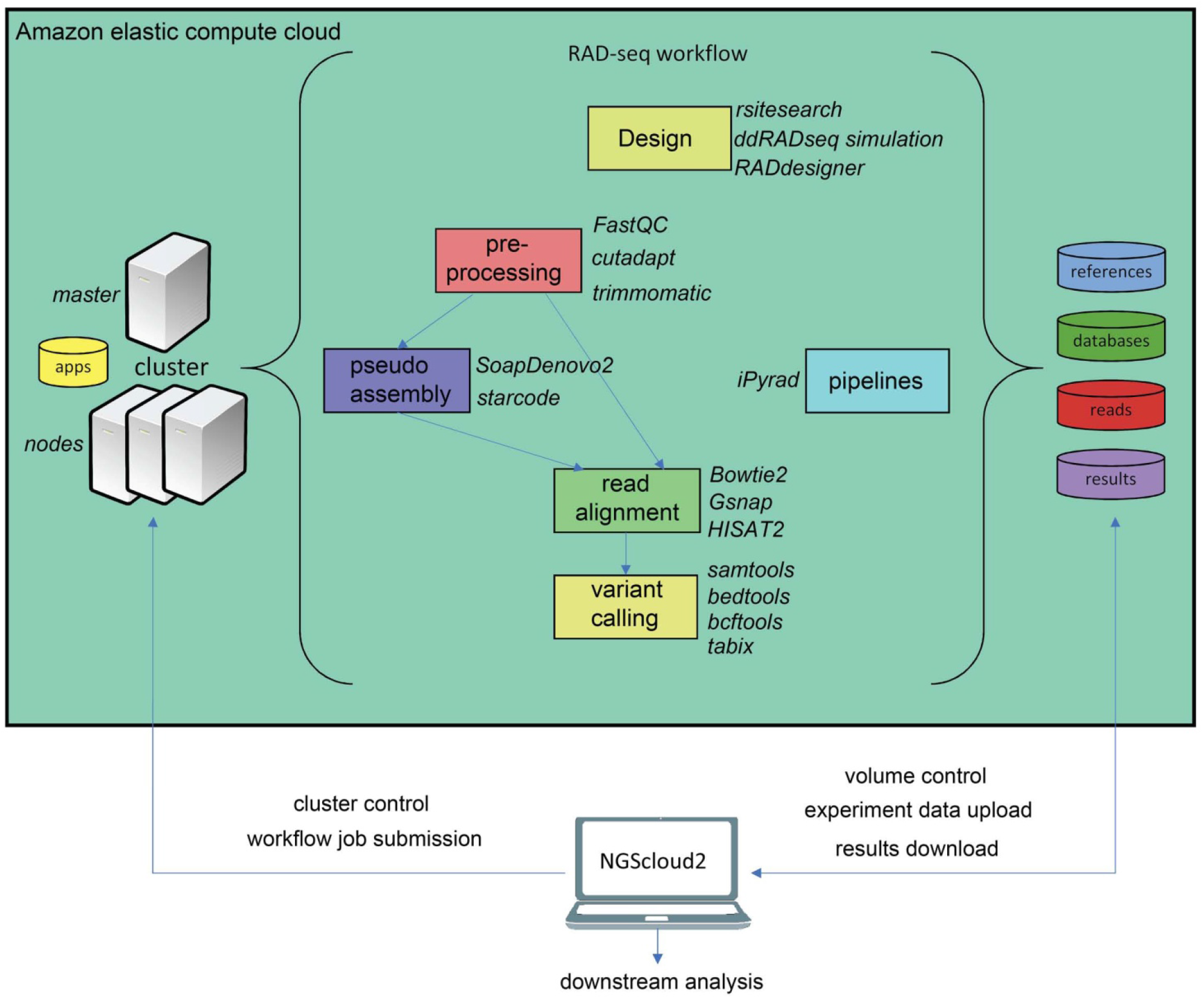
Reference-based RADseq workflow in NGScloud2.

#### Functional annotation

As a last improvement over the original version, NGScloud2 encapsulates our standalone application TOA (Mora-Márquez et al., 2020b), so it can run in EC2. This application automates the extraction of functional information from genomic databases, both plant specific (PLAZA) and general-purpose genomic databases (NCBI’s RefSeq and NR/NT), and the annotation of sequences (Figure 5). TOA can be a good complement for both RNAseq and ddRADseq workflows in non-model plant species that has shown optimal performance in AWS’s EC2 cloud. TOA aims to establish workflows geared towards woody plant species that automate the extraction of information from genomic databases and the annotation of sequences. TOA uses the following databases: Dicots PLAZA 4.0, Monocots PLAZA 4.0, Gymno PLAZA 1.0, NCBI RefSeq Plant and NCBI Nucleotide Database (NT) and NCBI Non-Redundant Protein Sequence Database (NR). Although TOA was primarily designed to work with woody plant species, it can also be used in the analysis of experiments on any type of plant organism. Additionally, NCBI Gene, InterPro and Gene Ontology databases are also used to complete the information.

**Figure 5.**
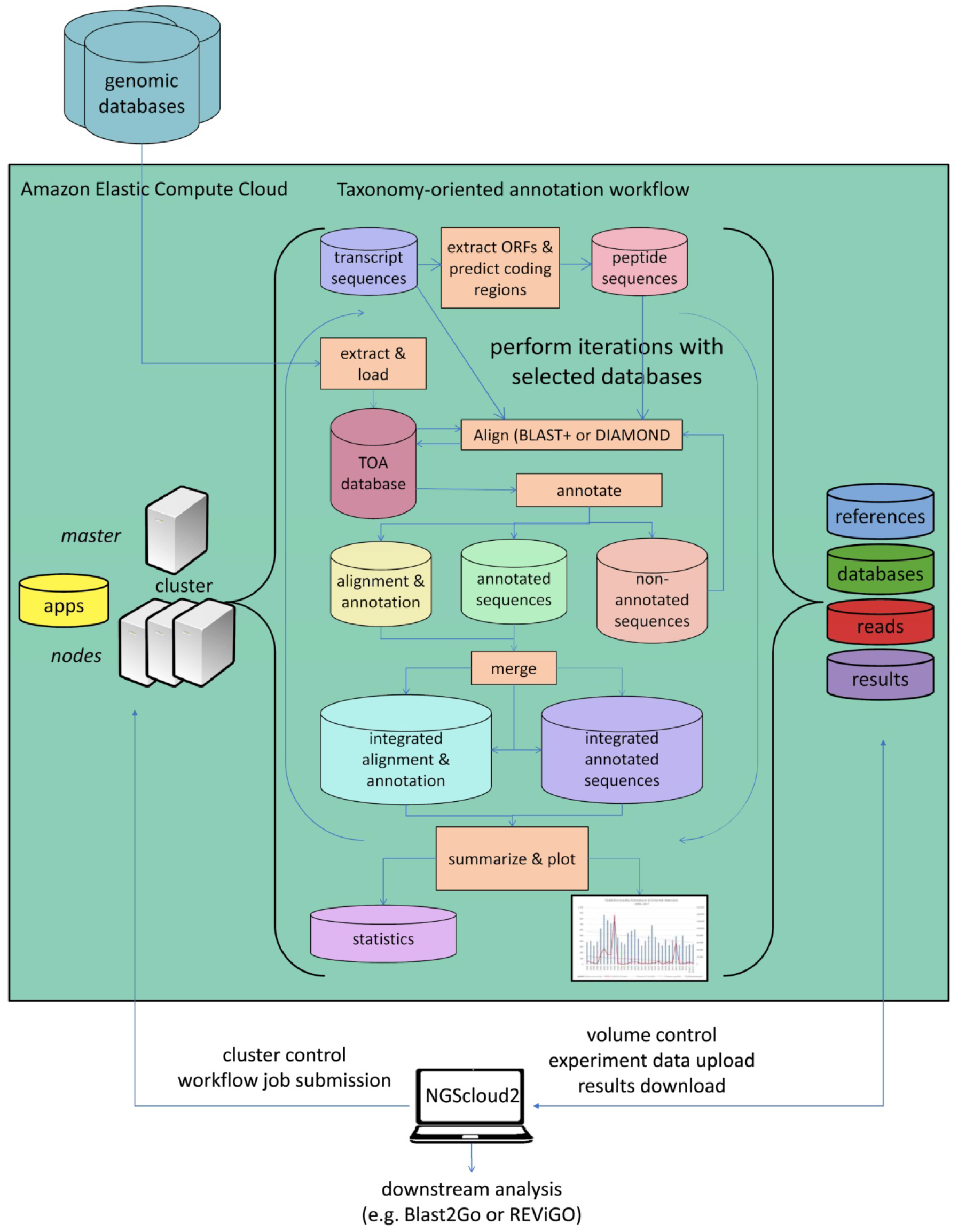
Functional annotation workflow in NGScloud2.

#### NGShelper

Besides the cloud infrastructure deployed in NGScloud2, we have included a companion package, NGShelper that contains python utilities to post-process the output of NGScloud2 pipelines. The package contains some Bash (Linux) and Bat (Windows) scripts to facilitate running the Python3 programs.

NGShelper facilitates format conversion of output files, filtering and subsetting of results, VCF and FASTA files statistics extraction, among others. Utilities list and their usage and parameters can be consulted at https://github.com/GGFHF/NGShelper/blob/master/Package/help.txt.

## Conclusions

NGScloud2 has significantly expanded the types of bioinformatic workflows to run using Amazon Web Services since its previous version. This new version has incorporated major technical improvements that optimize the use of popular software applications otherwise inaccessible to non-specialized users lacking suitable hardware infrastructures. Moreover, these technical improvements are oriented to significantly reduce costs by simplifying data access and taking advantage of EC2 spot instances that may produce savings of up to 50-80% in many steps of the analysis.

## Acknowledgements

This work was supported by the Spanish Ministry of Economy and Competitiveness-MINECO [grant number AGL2015-67495-C2-2-R]; the Spanish Ministry of Science and Innovation [grant numbers and PID2019-110330GB-C22 RTI2018-096465-B-I00]; the Regional Government of Madrid [grant numbers P2018/TCS4499 and S2018/TCS-4499]; and by an Amazon Research Grant. We’d like to thank M. Hurtado and Dr. V.M. Chano for beta-testing of NGScloud2 operability.

