## Supplemental Table 1 for "NGScloud2: optimized bioinformatic analysis using Amazon Web Services"

**Supplemental Table 1.** On-demand pricing and spot data corresponding to instances maneged by NGScloud2.
**OD**: On-demand price; **SoOD**: Saving over on-demand price (calcutaled over the thas 30 days); **FoI**: Frequency of interruption
Spot instace data published by AWS on on November 22th, 2020 at https://aws.amazon.com/ec2/spot/instance-advisor/ and on-demand pricing data published at https://aws.amazon.com/ec2/pricing/on-demand/

| Use | Type | Model | VCPU | Memory (GiB) | N. Virginia (us-east-1) | | | Paris (eu-west-3) | | | Generation |
|---|---|---|---|---|---|---|---|---|---|---|---|
| | | | | | OD ($/hour) | SoOD | FoI | OD ($/hour) | SoOD | FoI | |
| general purpose | t2 | t2.micro | 1 | 1,000 | 0,0116 | 70% | < 5% | 0,0132 | 70% | <5% | current |
| general purpose | t2 | t2.small | 1 | 2,000 | 0,0230 | 70% | < 5% | 0,0264 | 70% | 5-10% | current |
| general purpose | t2 | t2.medium | 2 | 4,000 | 0,0464 | 70% | 10-15% | 0,0528 | 70% | <5% | current |
| general purpose | t2 | t2.large | 2 | 8,000 | 0,0928 | 70% | >20% | 0,1056 | 70% | <5% | current |
| general purpose | t2 | t2.xlarge | 4 | 16,000 | 0,1856 | 70% | 5-10% | 0,2112 | 70% | 5-10% | current |
| general purpose | t2 | t2.2xlarge | 8 | 32,000 | 0,3712 | 70% | >20% | 0,4224 | 70% | <5% | current |
| general purpose | t3 | t3.micro | 2 | 1,000 | 0,0104 | 70% | < 5% | 0,0118 | 70% | 5-10% | current |
| general purpose | t3 | t3.small | 2 | 2,000 | 0,0208 | 70% | < 5% | 0,0236 | 70% | 5-10% | current |
| general purpose | t3 | t3.medium | 2 | 4,000 | 0,0416 | 70% | < 5% | 0,0472 | 70% | <5% | current |
| general purpose | t3 | t3.large | 2 | 8,000 | 0,0832 | 70% | 10-15% | 0,0944 | 70% | 10-15% | current |
| general purpose | t3 | t3.xlarge | 4 | 16,000 | 0,1664 | 70% | 15-20% | 0,1888 | 70% | >20% | current |
| general purpose | t3 | t3.2xlarge | 8 | 32,000 | 0,3328 | 70% | 5-10% | 0,3776 | 70% | 5-10% | current |
| general purpose | t3a | t3a.micro | 2 | 1,000 | 0,0094 | 70% | < 5% | 0,0106 | 70% | 5-10% | current |
| general purpose | t3a | t3a.small | 2 | 2,000 | 0,0188 | 70% | < 5% | 0,0212 | 70% | 10-15% | current |
| general purpose | t3a | t3a.medium | 2 | 4,000 | 0,0376 | 70% | < 5% | 0,0425 | 70% | 5-10% | current |
| general purpose | t3a | t3a.large | 2 | 8,000 | 0,0752 | 70% | < 5% | 0,0850 | 70% | 10-15% | current |
| general purpose | t3a | t3.xlarge | 4 | 16,000 | 0,1504 | 70% | < 5% | 0,1699 | 70% | 10-15% | current |
| general purpose | t3a | t3a.2xlarge | 8 | 32,000 | 0,3008 | 70% | < 5% | 0,3398 | 70% | <5% | current |
| general purpose | m3 | m3.medium | 1 | 3,750 | 0,0670 | 90% | 10-15% | N/A | N/A | N/A | previous |
| general purpose | m3 | m3.large | 2 | 7,500 | 0,1330 | 77% | < 5% | N/A | N/A | N/A | previous |
| general purpose | m3 | m3.xlarge | 4 | 15,000 | 0,2660 | 77% | < 5% | N/A | N/A | N/A | previous |
| general purpose | m3 | m3.2xlarge | 8 | 30,000 | 0,5320 | 76% | < 5% | N/A | N/A | N/A | previous |
| general purpose | m4 | m4.large | 2 | 8,000 | 0,1000 | 59% | < 5% | N/A | N/A | N/A | current |
| general purpose | m4 | m4.xlarge | 4 | 16,000 | 0,2000 | 62% | < 5% | N/A | N/A | N/A | current |
| general purpose | m4 | m4.2xlarge | 8 | 32,000 | 0,4000 | 57% | 5-10% | N/A | N/A | N/A | current |
| general purpose | m4 | m4.4xlarge | 16 | 64,000 | 0,8000 | 60% | >20% | N/A | N/A | N/A | current |
| general purpose | m4 | m4.10xlarge | 40 | 160,000 | 2,0000 | 67% | >20% | N/A | N/A | N/A | current |
| general purpose | m4 | m4.16xlarge | 64 | 256,000 | 3,2000 | 64% | 15-20% | N/A | N/A | N/A | current |
| general purpose | m5 | m5.large | 2 | 8,000 | 0,0960 | 56% | 10-15% | 0,1120 | 69% | 5-10% | current |
| general purpose | m5 | m5.xlarge | 4 | 16,000 | 0,1920 | 60% | 5-10% | 0,2240 | 72% | >20% | current |
| general purpose | m5 | m5.2xlarge | 8 | 32,000 | 0,3840 | 58% | 5-10% | 0,4480 | 71% | 15-20% | current |
| general purpose | m5 | m5.4xlarge | 16 | 64,000 | 0,7680 | 60% | 10-15% | 0,8960 | 72% | <5% | current |
| general purpose | m5 | m5.8xlarge | 32 | 128,000 | 1,5360 | 63% | >20% | 1,7920 | 72% | 15-20% | current |
| general purpose | m5 | m5.12xlarge | 48 | 192,000 | 2,3040 | 65% | >20% | 2,6880 | 72% | <5% | current |
| general purpose | m5 | m5.16xlarge | 64 | 256,000 | 3,0720 | 62% | 15-20% | 3,5840 | 72% | 15-20% | current |
| general purpose | m5 | m5.24xlarge | 96 | 384,000 | 4,6080 | 71% | 10-15% | 5,3760 | 72% | 5-10% | current |
| general purpose | m5a | m5a.large | 2 | 8,000 | 0,0860 | 54% | <5% | 0,1010 | 69% | 10-15% | current |
| general purpose | m5a | m5a.xlarge | 4 | 16,000 | 0,1720 | 55% | 10-15% | 0,2020 | 69% | 10-15% | current |
| general purpose | m5a | m5a.2xlarge | 8 | 32,000 | 0,3440 | 52% | 10-15% | 0,4040 | 68% | >20% | current |
| general purpose | m5a | m5a.4xlarge | 16 | 64,000 | 0,6880 | 55% | 10-15% | 0,8080 | 69% | 15-20% | current |
| general purpose | m5a | m5a.8xlarge | 32 | 128,000 | 1,3760 | 55% | 10-15% | 1,6160 | 69% | 10-15% | current |
| general purpose | m5a | m5a.12xlarge | 48 | 192,000 | 2,0640 | 59% | >20% | 2,4240 | 68% | 15-20% | current |
| general purpose | m5a | m5a.16xlarge | 64 | 256,000 | 2,7520 | 60% | >20% | 3,2320 | 68% | 15-20% | current |
| general purpose | m5a | m5a.24xlarge | 96 | 384,000 | 4,1280 | 60% | 15-20% | 4,8480 | 69% | 10-15% | current |
| compute optimized | c3 | c3.large | 2 | 3,750 | 0,1050 | 72% | <5% | N/A | N/A | N/A | previous |
| compute optimized | c3 | c3.xlarge | 4 | 7,500 | 0,2100 | 70% | <5% | N/A | N/A | N/A | previous |
| compute optimized | c3 | c3.2xlarge | 8 | 15,000 | 0,4200 | 62% | >20% | N/A | N/A | N/A | previous |
| compute optimized | c3 | c3.4xlarge | 16 | 30,000 | 0,8400 | 64% | >20% | N/A | N/A | N/A | previous |
| compute optimized | c3 | c3.8xlarge | 32 | 60,000 | 1,6800 | 70% | >20% | N/A | N/A | N/A | previous |

| | | | | | | | | | | | |
|---|---|---|---|---|---|---|---|---|---|---|---|
| compute optimized | c4 | c4.large | 2 | 3,750 | 0,1000 | 67% | 5-10% | N/A | N/A | N/A | current |
| compute optimized | c4 | c4.xlarge | 4 | 7,500 | 0,1990 | 63% | 15-20% | N/A | N/A | N/A | current |
| compute optimized | c4 | c4.2xlarge | 8 | 15,000 | 0,3980 | 57% | 5-10% | N/A | N/A | N/A | current |
| compute optimized | c4 | c4.4xlarge | 16 | 30,000 | 0,7960 | 69% | >20% | N/A | N/A | N/A | current |
| compute optimized | c4 | c4.8xlarge | 36 | 60,000 | 1,5910 | 66% | >20% | N/A | N/A | N/A | current |
| compute optimized | c5 | c5.large | 2 | 4,000 | 0,0850 | 62% | <5% | 0,1010 | 70% | 5-10% | current |
| compute optimized | c5 | c5.xlarge | 4 | 8,000 | 0,1700 | 57% | <5% | 0,2020 | 70% | 5-10% | current |
| compute optimized | c5 | c5.2xlarge | 8 | 16,000 | 0,3400 | 57% | <5% | 0,4040 | 59% | >20% | current |
| compute optimized | c5 | c5.4xlarge | 16 | 32,000 | 0,6800 | 60% | 10-15% | 0,8080 | 70% | <5% | current |
| compute optimized | c5 | c5.9xlarge | 36 | 72,000 | 1,5300 | 61% | 10-15% | 1,8180 | 70% | 5-10% | current |
| compute optimized | c5 | c5.12xlarge | 48 | 96,000 | 2,0400 | 62% | 10-15% | 2,4240 | 70% | 10-15% | current |
| compute optimized | c5 | c5.18xlarge | 72 | 144,000 | 3,0600 | 62% | <5% | 3,6360 | 70% | <5% | current |
| compute optimized | c5 | c5.24xlarge | 96 | 192,000 | 4,0800 | 62% | <5% | 4,8480 | 70% | 10-15% | current |
| compute optimized | c5a | c5a.large | 2 | 4,000 | 0,0770 | 57% | 5-10% | N/A | N/A | N/A | current |
| compute optimized | c5a | c5a.xlarge | 4 | 8,000 | 0,1540 | 57% | 10-15% | N/A | N/A | N/A | current |
| compute optimized | c5a | c5a.2xlarge | 8 | 16,000 | 0,3080 | 57% | <5% | N/A | N/A | N/A | current |
| compute optimized | c5a | c5a.4xlarge | 16 | 32,000 | 0,6160 | 51% | 10-15% | N/A | N/A | N/A | current |
| compute optimized | c5a | c5a.8xlarge | 32 | 64,000 | 1,2320 | 58% | 10-15% | N/A | N/A | N/A | current |
| compute optimized | c5a | c5a.12xlarge | 48 | 96,000 | 1,8480 | 47% | 5-10% | N/A | N/A | N/A | current |
| compute optimized | c5a | c5a.16xlarge | 64 | 128,000 | 2,4640 | 58% | 10-15% | N/A | N/A | N/A | current |
| compute optimized | c5a | c5a.24xlarge | 96 | 192,000 | 3,6960 | 58% | 10-15% | N/A | N/A | N/A | current |
| memory optimized | r3 | r3.large | 2 | 3,750 | 0,1660 | 80% | <5% | N/A | N/A | N/A | previous |
| memory optimized | r3 | r3.xlarge | 4 | 30,500 | 0,3330 | 79% | <5% | N/A | N/A | N/A | previous |
| memory optimized | r3 | r3.2xlarge | 8 | 61,000 | 0,6650 | 79% | 5-10% | N/A | N/A | N/A | previous |
| memory optimized | r3 | r3.4xlarge | 16 | 122,000 | 1,3300 | 79% | 5-10% | N/A | N/A | N/A | previous |
| memory optimized | r3 | r3.8xlarge | 32 | 244,000 | 2,6600 | 80% | 10-15% | N/A | N/A | N/A | previous |
| memory optimized | r4 | r4.large | 2 | 15,250 | 0,1330 | 74% | 5-10% | 0,1560 | 80% | <5% | current |
| memory optimized | r4 | r4.xlarge | 4 | 30,500 | 0,2660 | 65% | <5% | 0,3120 | 80% | 5-10% | current |
| memory optimized | r4 | r4.2xlarge | 8 | 61,000 | 0,5320 | 67% | <5% | 0,6240 | 80% | <5% | current |
| memory optimized | r4 | r4.4xlarge | 16 | 122,000 | 1,0640 | 74% | 10-15% | 1,2480 | 80% | >20% | current |
| memory optimized | r4 | r4.8xlarge | 32 | 244,000 | 2,1280 | 74% | 10-15% | 2,4960 | 80% | 10-15% | current |
| memory optimized | r4 | r4.16xlarge | 64 | 488,000 | 4,2560 | 74% | 10-15% | 4,9920 | 80% | 10-15% | current |
| memory optimized | r5 | r5.large | 2 | 16,000 | 0,1260 | 72% | <5% | 0,1480 | 77% | >20% | current |
| memory optimized | r5 | r5.xlarge | 4 | 32,000 | 0,2520 | 68% | <5% | 0,2960 | 71% | >20% | current |
| memory optimized | r5 | r5.2xlarge | 8 | 64,000 | 0,5040 | 65% | 5-10% | 0,5920 | 77% | 5-10% | current |
| memory optimized | r5 | r5.4xlarge | 16 | 128,000 | 1,0080 | 68% | <5% | 1,1840 | 77% | 15-20% | current |
| memory optimized | r5 | r5.8xlarge | 32 | 256,000 | 2,0160 | 71% | 5-10% | 2,3680 | 77% | <5% | current |
| memory optimized | r5 | r5.12xlarge | 48 | 384,000 | 3,0240 | 72% | 5-10% | 3,5520 | 77% | 10-15% | current |
| memory optimized | r5 | r5.16xlarge | 64 | 512,000 | 4,0320 | 71% | <5% | 4,7360 | 0% | <5% | current |
| memory optimized | r5 | r5.24xlarge | 96 | 768,000 | 6,0480 | 72% | <5% | 7,1040 | 77% | 10-15% | current |
| memory optimized | r5a | r5a.large | 2 | 16,000 | 0,1130 | 68% | 5-10% | 0,1330 | 75% | 10-15% | current |
| memory optimized | r5a | r5a.xlarge | 4 | 32,000 | 0,2260 | 60% | 5-10% | 0,2660 | 75% | 10-15% | current |
| memory optimized | r5a | r5a.2xlarge | 8 | 64,000 | 0,4520 | 68% | 10-15% | 0,5320 | 75% | 5-10% | current |
| memory optimized | r5a | r5a.4xlarge | 16 | 128,000 | 0,9040 | 66% | 10-15% | 1,0640 | 75% | 10-15% | current |
| memory optimized | r5a | r5a.8xlarge | 32 | 256,000 | 1,8080 | 68% | >20% | 2,1280 | 75% | 5-10% | current |
| memory optimized | r5a | r5a.12xlarge | 48 | 384,000 | 2,7120 | 63% | >20% | 3,1920 | 75% | 10-15% | current |
| memory optimized | r5a | r5a.16xlarge | 64 | 512,000 | 3,6160 | 68% | 10-15% | 4,2560 | 75% | 10-15% | current |
| memory optimized | r5a | r5a.24xlarge | 96 | 768,000 | 5,4240 | 68% | 15-20% | 6,3840 | 75% | 10-15% | current |
