## Supplementary material for "NGScloud2: optimized bioinformatic analysis using Amazon Web Services": Table 2

**Supplemental Table 2.** Processes run in *De novo* RNA-seq tests.
Benchmark read data: paired read files of library Pcan1-S0 included in NCBI BioProject with accession PRJNA510935.

| NGScloud2 menu item | Comment | Result dataset id | Instance | vCPU | Memory (GiB) | Purchasing op | Elapsed time (s) |
|---|---|---|---|---|---|---|---|
| Datasets > Read dataset file transfer | Compressed Pcan FASTQ files (*) | local process | t3.medium | 2 | 4 | spot | - |
| De novo RNA-seq > Read quality > FastQC | - | fastqc-201126-185600 | t3.medium | 2 | 4 | spot | 500 |
| De novo RNA-seq > Trimming > Trimmomatic | Cut 12 nucleotides from the start of the read | trimmo-201126-190912 | t3.medium | 2 | 4 | spot | 1.474 |
| De novo RNA-seq > Read quality > FastQC | - | fastqc-201126-193927 | t3.medium | 2 | 4 | spot | 407 |
| Datasets > Read dataset file compression/decompression | Trimmed reads | gzip-201126-195230 | t3.medium | 2 | 4 | spot | 259 |
| De novo RNA-seq > Assembly > SOAPdenovo-Trans | Trimmed reads | sdnt-201126-202511 | r5a.2xlarge | 8 | 64 | spot | 2.486 |
| De novo RNA-seq > Read alignment > Bowtie2 | Trimmed reads with assembled transcriptome | bowtie2-201126-213009 | r5a.2xlarge | 8 | 64 | spot | 2.062 |
| De novo RNA-seq > Transcriptome quelity assessment > BUSCO | Assembled transcriptome | busco-201128-140210 | r5a.2xlarge | 8 | 64 | spot | 629 |
| De novo RNA-seq > Transcriptome quelity assessment > QUAST | Assembled transcriptome | quast-201127-181647 | r5a.xlarge | 4 | 32 | spot | 15 |
| De novo RNA-seq > Transcriptome quelity assessment > rnaQUAST | Assembled transcriptome and trimmed reads | rnaquast-201130-104048 | r5a.2xlarge | 8 | 64 | spot | 1.948 |
| De novo RNA-seq > Transcriptome quelity assessment > RSEM-EVAL | Assembled transcriptome and trimmed reads | rsemeval-201129-152226 | r5a.2xlarge | 8 | 64 | spot | 1.979 |
| De novo RNA-seq > Transcriptome quelity assessment > Transrate | Assembled transcriptome and trimmed reads | transrate-201128-000833 | r5a.xlarge | 4 | 32 | spot | 2.036 |
| De novo RNA-seq > Transcriptome filtering > CD-HIT-EST | Assembled transcriptome | cdhitest-201129-093523 | r5a.xlarge | 4 | 32 | spot | 15.952 |
| De novo RNA-seq > Transcriptome filtering > transcript-filter | dataset rsemeval-201129-152226 | transfil-201129-185702 | t3.medium | 2 | 4 | spot | 7 |
| De novo RNA-seq > Quantitation > eXpress | Assembled transcriptome and dataset bowtie2-201126-213009 | express-201129-092047 | r5a.xlarge | 4 | 32 | spot | 320 |
| De novo RNA-seq > Quantitation > kallisto | Assembled transcriptome and trimmed reads | kallisto-201129-085931 | r5a.xlarge | 4 | 32 | spot | 1.168 |
| De novo RNA-seq > Variant calling > Variant calling | Assembled transcriptome and dataset bowtie2-201126-213009 | varcalling-201129-163739 | r5a.2xlarge | 8 | 64 | spot | 4.641 |
