## Supplementary material for "NGScloud2: optimized bioinformatic analysis using Amazon Web Services": Table 3

**Supplemental Table 3.** Processes run in Reference-based RNA-seq tests.
Benchmark read data: paired read files of 23 libraries included in NCBI BioProject with accession PRJNA510935.

| NGScloud2 menu item | Comment | Result dataset id | Instance | vCPU | Memory (GiB) | Purchasing option | Elapsed time (s) |
|---|---|---|---|---|---|---|---|
| Datasets > Reference dataset file transfer | Compressed genome and annotation files of Pinus taeda | local process | t3.medium | 2 | 4 | spot | - |
| Datasets > Reference dataset file compression/decompression | Compressed genome and annotation files of P. taeda | gzip-201104-103050 | t3.medium | 2 | 4 | spot | 334 |
| Datasets > Read dataset file transfer | Compressed Pcan FASTQ files | local process | t3.medium | 2 | 4 | spot | - |
| Reference-based RNA-seq > Read quality > FastQC | - | fastqc-201103-144624 | t3.medium | 2 | 4 | spot | 15.642 |
| Reference-based RNA-seq > Trimming > Trimmomatic | Cut 12 nucleotides from the start of the read | trimmo-201103-212135 | t3.medium | 2 | 4 | spot | 35.758 |
| Reference-based RNA-seq > Read quality > FastQC | - | fastqc-201103-144624 | t3.medium | 2 | 4 | spot | 11.631 |
| Datasets > Read dataset file compression/decompression | Trimmed reads | gzip-201104-144601 | t3.medium | 2 | 4 | spot | 9.169 |
| Reference-based RNA-seq > Read alignment > HISAT2 | Trimmed reads with P. taeda genome | hisat2-20117-131003 | r5.8xlarge | 32 | 256 | on-demand | 73.516 |
| Reference-based RNA-seq > Quantitation > htseq-count | HISAT2 read alignment and P. taeda annotation file | htseqcount-201121-125750 | m5.2xlarge | 8 | 32 | spot | 9.113 |
| Reference-based RNA-seq > Variant calling > Variant calling | HISAT2 read alignment and P. Taeda genome | varcalling-201123-133906 | m5.2xlarge | 8 | 32 | spot | 60.180 |
| Reference-based RNA-seq > Transcriptome alignment > GMAP | Pinus canariensis transcriptome with P. Taeda genome | gmap-201118-122401 | r5.8xlarge | 32 | 256 | on-demand | 35.039 |
