## Supplementary material for "NGScloud2: optimized bioinformatic analysis using Amazon Web Services": Table 4

**Supplemental Table 4.** Processes run in Tree-oriented Annotation pipeline tests.
Benchmark read data: SRR8199746_2.fastq.gz, SRR8199747_2.fastq.gz, SRR8199748_2.fastq.gz, SRR8199749_2.fastq.gz, SRR8199750_2.fastq.gz, SRR8199751_2.fastq.gz, SRR8199760_2.fastq.gz and SRR8199761_2.fastq.gz. These files are included in NCBI BioProject with accession PRJNA505763.

| NGScloud2 menu item | Comment | Result dataset id | Instance | vCPU | Memory (GiB) | Purchasing option | Elapsed time (s) |
|---|---|---|---|---|---|---|---|
| Datasets > Read dataset file transfer | Upload compressed Qilex x Qsuber FASTQ files | local process | t3.medium | 2 | 4 | spot | - |
| RAD-seq > Read quality > FastQC | Study read quality | fastqc-201028-124325 | t3.medium | 2 | 4 | spot | 357 |
| RAD-seq > Trimming > Trimmomatic | Cut restiction site sequences | trimmo-201028-131122 | t3.medium | 2 | 4 | spot | 2.241 |
| RAD-seq > Read quality > FastQC | Study read quality | fastqc-201028-124325 | t3.medium | 2 | 4 | spot | 357 |
| RAD-seq > Trimming > cutadapt | Remove Illumina TruSeq adapter sequences | cutadapt-201028-154815 | t3.medium | 2 | 4 | spot | 1.589 |
| RAD-seq > Read quality > FastQC | Study read quality | fastqc-201028-161935 | t3.medium | 2 | 4 | spot | 342 |
| RAD-seq > Trimming > cutadapt | Remove overrepresented sequences | cutadapt-201028-184743 | t3.medium | 2 | 4 | spot | 1.441 |
| RAD-seq > Read quality > FastQC | Study read quality | fastqc-201028-191549 | t3.medium | 2 | 4 | spot | 331 |
| RAD-seq > Trimming > cutadapt | Remove poly(A) tail sequences | cutadapt-201029-142345 | t3.medium | 2 | 4 | spot | 1.321 |
| RAD-seq > Read quality > FastQC | Study read quality | fastqc-201029-150630 | t3.medium | 2 | 4 | spot | 362 |
| Datasets > Read dataset file compression/decompre | Decompress final trimmed reads | gzip-201029-152047 | t3.medium | 2 | 4 | spot | 100 |
| RAD-seq > Pseudo assembly > starcode | Pseudo assemble reads | starcode-201029-161928 | r5.2xlarge | 8 | 64 | on-demand | 13.874 |
| RAD-seq > Pseudo assembly > SOAPdenovo2 | Pseudo assemble reads | sdn2-201029-201719 | r5.2xlarge | 8 | 64 | on-demand | 1.907 |
| RAD-seq > Read alignment > Bowtie2 | Align reads to starcode pseudo assembly (alignment 1) | bowtie2-201030-130308 | r5.2xlarge | 8 | 64 | on-demand | 3.355 |
| RAD-seq > Read alignment > Bowtie2 | Align reads to SOAPdenovo2 pseudo assembly (alignment 2) | bowtie2-201030-205451 | r5.2xlarge | 8 | 64 | on-demand | 1.702 |
| RAD-seq > Read alignment > GSNAP | Align reads to starcode pseudo assembly (alignment 3) | gsnap-201030-124300 | r5.2xlarge | 8 | 64 | on-demand | 954 |
| RAD-seq > Read alignment > GSNAP | Align reads to SOAPdenovo2 pseudo assembly (alignment 4) | gsnap-201030-144844 | r5.2xlarge | 8 | 64 | on-demand | 1.672 |
| RAD-seq > Variant calling > Variant calling | Search variants using starcode pseudo assembly and alignment 1 | varcalling-201101-175723 | r5.2xlarge | 8 | 64 | spot | 3.171 |
| RAD-seq > Variant calling > Variant calling | Search variants using starcode pseudo assembly and alignment 3 | varcalling-201031-141134 | r5.2xlarge | 8 | 64 | spot | 3.152 |
| RAD-seq > Variant calling > Variant calling | Search variants using SOAPdenovo2 pseudo assembly and alignment 2 | varcalling-201101-143346 | r5.2xlarge | 8 | 64 | spot | 532 |
| RAD-seq > Variant calling > Variant calling | Search variants using SOAPdenovo2 pseudo assembly and alignment 4 | varcalling-201031-153253 | r5.2xlarge | 8 | 64 | spot | 3.863 |
