## Supplementary material for "NGScloud2: optimized bioinformatic analysis using Amazon Web Services": Table 5

**Supplemental Table 5.** Processes run in Tree-oriented Annotation pipeline tests.
Benchmark read data: MonolignolsGenes.fasta. Raes J, Rohde A, Christensen JH, Van de Peer Y, Boerjan W. 2003. Genome-Wide Characterization of the Lignification Toolbox in Arabidopsis. Plant Physiology 133:1051–1071. DOI: 10.1104/pp.103.026484.

| NGScloud2 menu item | Comment | Result dataset id | Instance | vCPU | Memory (GiB) | Purchasing option | Elapsed time (s) |
|---|---|---|---|---|---|---|---|
| Datasets > Reference dataset file transfer | File MonolignolsGenes.fasta | local process | t3.medium | 2 | 4 | spot | - |
| Taxonomy-oriented annotation > Annotation pipelines > TOA nucleotide pipeline | Alignment tool: BLAST+ | toapipelinent-201125-211743 | r5.xlarge | 4 | 32 | spot | 1.041 |
| Taxonomy-oriented annotation > Annotation pipelines > TOA nucleotide pipeline | Alignment tool: DIAMOND | toapipelinent-201125-213659 | r5.xlarge | 4 | 32 | spot | 166 |
| Taxonomy-oriented annotation > Annotation pipelines > TOA amino acid pipeline | Alignment tool: BLAST+ | toapipelineaa-201125-214039 | r5.xlarge | 4 | 32 | spot | 4.997 |
| Taxonomy-oriented annotation > Annotation pipelines > TOA amino acid pipeline | Alignment tool: DIAMOND | toapipelineaa-201125-231005 | r5.xlarge | 4 | 32 | spot | 7.006 |
| Taxonomy-oriented annotation > Annotation pipelines > Annotation merger of TOA pipelines | Merger of toapipelinent-201125-211743 & toapipelineaa-201125-214039 | toamergeann-201126-114035 | t3.medium | 2 | 4 | spot | 7 |
| Taxonomy-oriented annotation > Annotation pipelines > Annotation merger of TOA pipelines | Merger of toapipelinent-201125-213659 & toapipelineaa-201125-231005 | toamergeann-201126-114918 | t3.medium | 2 | 4 | spot | 1 |
